# Alkalinity modulates a unique suite of genes to recalibrate growth and pH homeostasis

**DOI:** 10.1101/2022.12.12.520164

**Authors:** Mitylene Bailey, En-Jung Hsieh, Huei-Hsuan Tsai, Arya Ravindran, Wolfgang Schmidt

## Abstract

Alkaline soils pose a conglomerate of constraints to plants, restricting the growth and fitness of non-adapted species in habitats with low active proton concentrations. To thrive under such conditions, plants have to compensate for a potential increase in cytosolic pH and restricted softening of the cell wall to invigorate cell elongation in a proton-depleted environment. To discern mechanisms that aid in the adaptation to external pH, we grew plants on media with pH values ranging from 4.5 to 8.5. Growth was severely restricted at pH 4.5 and above pH 6.5, and associated with decreasing chlorophyll levels at alkaline pH. Bicarbonate treatment worsened plant performance, suggesting effects that differ from those exerted by pH as such. Transcriptional profiling of roots subjected to short-term transfer from optimal (pH 5.5) to alkaline (pH 7.5) media unveiled a large set of differentially expressed genes that were partially congruent with genes affected by low pH, bicarbonate and nitrate, but showed only a very small overlap with genes responsive to the availability of iron. Further analysis of selected genes disclosed pronounced responsiveness of their expression over a wide range of external pH values. Alkalinity altered the expression of various proton/anion co-transporters, possibly to recalibrate cellular proton homeostasis. Co-expression analysis of pH-responsive genes identified a module of genes encoding proteins with putative functions in the regulation of root growth, which appears to be conserved in plants subjected to low pH or bicarbonate. Our analysis provides an inventory of pH-sensitive genes and allows comprehensive insights into processes that are orchestrated by external pH.

## Introduction

Soil pH, a composite readout derived from weathering, climate, vegetation, and the parental material, exerts a multitude of effects on the plant’s productivity and fitness. While the concentration of hydrogen ions can vary both spatially and temporally, the pH of the soil solution is characteristic of a certain habitat and soil type, and governs the composition of natural plant communities by defining the availability of nutrients, the activity of soil microbial assemblages, the seed germination rate, and the available water capacity (Tsai and Schmidt, 2021). Soil pH can range from extreme acidic (~pH 3) to extreme alkaline (~pH 9) conditions. In acid soils, the presence Al^3+^, released from clay minerals at a pH below 5 is the main factor that restricts species with insufficient detoxification mechanisms to thrive under such conditions. Alkaline soils, on the other hand, dramatically limit the availability of iron and phosphate, excluding so-called calcifuge (chalk-fleeing) species from these habitats. Alkaline calcareous soils contain high levels of calcium or magnesium carbonate that both increase and buffer soil pH, posing a further constraint on plants growing on such sites. Calcicole (chalk-loving) species that can tolerate these conditions are able to efficiently mine iron and phosphate from recalcitrant pools (Grime and Hodgson 1968). If - or to what extent - pH as such plays a role in defining calcicole behavior is not known.

Maintaining cytosolic pH within a narrow, slightly alkaline set value is critical to cell viability. To recalibrate intracellular proton concentration, plant cells employ a mechanism referred to as the biochemical pH stat (Smith and Raven, 1979), which depends on the production or consumption of protons during the carboxylation and decarboxylation of organic acids. In addition, electrogenic transport processes driven by plasma membrane-localized P-type and vacuolar V-ATPases contribute to cellular pH homeostasis, a regulatory mechanism referred to as biophysical pH stat. Migration of protons across the plasma membrane alters, however, both the cytoplasmic and apoplastic pH. In plants, an acidic apoplast is a prerequisite for loosening the cell wall to aid cell elongation (acid growth theory; Hager et al., 1971). Crucially, changing apoplastic pH – either by manipulating electrogenic proton fluxes across the plasma membrane or by exposing plants to media of different pH - alters root growth according to the prevailing hydrogen concentration (Li et al., 2021; Lin et al., 2021), corroborating this concept.

If and how plants sense the pH of the apoplast was a long-standing enigma (Tsai and Schmidt, 2021). Recently, a bimodal apoplastic pH sensing system was shown to negotiate the balance of two basic tasks of root cells, growth and defense (Liu et al., 2022a). In this system, a sulfotyrosine residue in the root growth factor (RGF) peptide is protonated at low pH, which facilitates binding to its receptor RGFR and meristem development by regulating PLETHORA protein gradients (Liu et al., 2022a; Aida et al., 2004). At elevated pH this residue is deprotonated, which compromises the formation of the RGF-RGFR complex. Alkalization of the apoplast engages a second component of the pH sensing system by removing protons from essential Glu and Asp residues of the receptor protein PEPR, which supports binding of its ligand PEP1 and triggers immunity.

Whether the RGF/PEP-based sensing system is also critical to the adaptation of plants to alkaline conditions remains to be elucidated. To shed light on the mechanisms that allow perception of external pH and pH-dependent adjustment of metabolism and growth, we conducted gene expression analyses of (calcifuge) *Arabidopsis* plants subjected to alkaline conditions. In addition, we analyzed the growth of plants exposed to media differing in pH, including a treatment where external pH was controlled by supplementing the growth media with bicarbonate. Our analysis revealed that short-term exposure to high pH engages or repress signaling cascades and transport processes that adapt plants to alkaline conditions. We further show that gene expression is modulated over a wide range of pH values, suggesting that a core set of pH-dependent genes is adapting plants to both acidic and alkaline soil pH. Transcriptomic analysis and a comparison with related transcriptomic studies disclosed modules with putative roles in intracellular pH homeostasis and root development. Root growth appears to be governed by a pH-dependent, redox-based mechanism that is strongly repressed by elevated pH.

## Materials and methods

### Pant growth conditions

Seeds of Arabidopsis (*A. thaliana* (L.) Heynh) accession Col-0 were obtained from the Arabidopsis Biological Resource Center (Ohio State University). Seeds were sterilized with 35% (v/v) bleach, washed five times with sterile water. Plants were grown in a growth chamber on solid nutrient media as described by Estelle and Somerville (1987) (ES media), composed of 5□mM KNO_3_, 2□mM MgSO_4_, 2□mM Ca(NO_3_)_2_, 2.5□mM KH_2_PO_4_, 70□μmM H_3_BO_3_, 14□μM MnCl_2_, 1□μM ZnSO_4_, 0.5 μM CuSO_4_, 0.01□μM CoCl_2_, 0.2□μM Na_2_MoO_4_, 1% (w/v) MES, and 1.5% (w/v) sucrose, solidified with 0.4% Gelrite pure (Kelco) and supplemented with 40□μM FeEDDHA. Plants were grown at 22°C under continuous light, 100 μmol m^−2^s^−1^ until time of collection. The pH was adjusted to various pH values as indicated. Media with a pH above 7.0 were buffered with 1g/L MOPS; for pH 4.5 to 6.5 media 1g/L MES was used as buffering agent.

### pH treatments

Ten-day-old plants were transferred from ES media pH 5.5 to plates containing media adjusted to pH 4.5 to 8.5 in increments of one unit. Additionally, plants were grown on plates supplemented with 3 mM NaHCO_3_ (pH ~6.7). Roots exposed for 6 hours or 14 days to the varying pH conditions were isolated from the shoots and flash-frozen in liquid nitrogen.

### RNA-seq analysis and definition of DEGs

Total RNA was isolated from roots of 10-day-old plants grown at optimal pH (5.5) or exposed for six-hour to pH 7.5 media using the RNeasy Plant Mini Kit (Qiagen). Three independent biological replicates were performed. Libraries for RNA-seq were prepared using the Illumina TruSeq RNA library Prep Kit (RS-122-2001, Illumina) according to the manufacturer’s protocol. Four μg of total RNA were used for library construction. PolyA RNA was captured by oligodT beads and fragmented when eluted from the beads. First-strand cDNA was synthesized from fragmented RNA using reverse transcriptase (SuperScrip III, Cat. No. 18080-093, Invitrogen) and random primers. The cDNA was then converted into double stranded DNA using the reagents supplied with the kit. Reactions were cleaned up with Agencourt AMPure XP beads (Beckman Coulter Genomics). Libraries were end-repaired, adenylated at the 3’ end, ligated with adapters, and amplified following the TruSeq™ RNA Sample Preparation v2 LS protocol. Finally, the products were purified and enriched with 10 cycles of PCR to create the final double-stranded cDNA library. For quality check, we used the BioRad QX200 Droplet Digital PCR EvaGreen supermix system (Cat. No. #1864034; BioRad, USA) and the Agilent High Sensitivity NGS Fragment Kit (1-6000 bp) (Cat. No. DNF-474-500; Agilent, USA). The prepared library was pooled for paired-end sequencing using Illumina the Novaseq 6000 at Taiwan Genome Industry Alliance Inc. (Taipei City, Taiwan) with 150 bp paired-ended sequence reads. RNAseq reads were pre-processed by trimmomatic (Bolger *et al*. 2014) for adapter removal (parameter: ILLUMINACLIP:TruSeq3-PE-2.fa:2:30:10:8:true MINLEN:80), where reads shorter than 80 bps were dropped. Passed reads were first mapped to transcript sequences of the Araport11 database using Bowtie2 (Langmead and Salzberg, 2012); only alignments of read pairs were accepted. The remaining reads were mapped to the Arabidopsis genome annotation (TAIR10) using BLAT (Kent, 2002). Read counts of genes were computed using the RackJ software package (https://rackj.sourceforge.net/) and normalized into log-count-per-million values using the TMM method (Robinson and Oshlack, 2010). Normalized RPKM values were inferred from the normalized log-count-per-million values. DEGs (differentially expressed genes) was identified as differentially expressed if the corresponding *P*-value was less than or equal to 0.05 and the fold-change was greater than two.

### RT-qPCR

Frozen plant samples were ground to powder using a Qiagen Tissue Lyser II for three cycles of 30 secs. RNA extraction was performed using TRIzol^®^ reagent; RNA was measured using a NanoDrop ND-1000 spectrophotometer. First-strand cDNA was synthesized using 1 μg RNA with the TOOLS Easy Fast RT Kit (Biotools). Three replicates of each sample measuring 10 μL reaction mixture made with AB SYBR Green PCR Master Mix (Applied Biosystems, Waltham, MA, USA) were used in the AB QuantStudio ™ 12K Flex Real Time PCR System (Applied Biosystems) following a preset program. Transcripts were measured using the comparative C_T_ (ΔΔC_T_) calculation method using multiple controls (EF1α and Tubulin). Graphs representing relative Expression (2^−ΔΔ*CT*^) were built using GraphPad Prism 9 for Windows 64-bit. Primers used for RT-qPCR are listed in Supplemental Table 1.

### Data analysis

Analysis of the Chen et al. 2021 data set was made by making comparisons of treated versus control samples using the Student’s t-test with an adjusted P-value. This, and all the other data were filtered using a log2f old-change of |1|. Venn diagrams were assembled using Venn diagram <http://bioinformatics.psb.ugent.be/webtools/Venn/>; heatmaps were designed by TBTools (Chen et al., 2020) both were edited in Inkscape.

### Co-expression analysis

To infer root-specific co-expression networks, a total of 1,194 root-specific RNA-seq data sets was downloaded from the Sequence Read Archive hosted by the National Center for Biotechnology Information and normalized. Networks were constructed with the MACCU toolbox (Lin et al., 2011) based on pair-wise comparison of the co-expression relationships of Arabidopsis genes expressed in using Pearson coefficients as indicated in the figures. Co-expression networks were edited in Cytoscape (Shannon et al., 2003).

### Plant growth characteristics

#### Fresh Weight, Rosette size, Chlorophyll Quantification

Whole shoots from 10- or 14-day-old plants grown on ES media with pH values ranging from pH 4.5 to 8.5 in increments of one or on media supplemented with NaHCO_3_ were weighed and chlorophyll content was measured using extractions made with 80% acetone modified from the protocol described by Lichtenthaler (1987). Rosette diameter was determined using the Image J measure tool (Schindelin et al., 2012).

## Results

### Growth of *Arabidopsis* plants is strongly affected by media pH

To gain a better understanding regarding the responses of *Arabidopsis* plants to changes in external pH, we grew plants for a two-week period on media adjusted to pH values from 4.5 to 8.5. In addition, we added a treatment in which the medium was supplemented with 3 mM bicarbonate, resulting in a media pH of 6.7. As expected from an accession that is not derived from or adapted to calcareous or alkaline condition, plants grew best on slightly acidic media with pH values of 5.5 or 6.5 (Fig. 1). Growth on media with strongly acidic pH (4.5) caused a steep decline in root and shoot weight; increasing the media pH to pH 7.5 or 8.5 dramatically decreased growth and chlorophyll concentration of the plants (Fig. 1). Bicarbonate-treated plants showed significant reduction in root and shoot weight, rosette diameter, and shoot-root ratio when compared to plants grown at comparable pH (6.5), indicative of effects that are independent of pH (Fig. 1).

**Figure 1.**
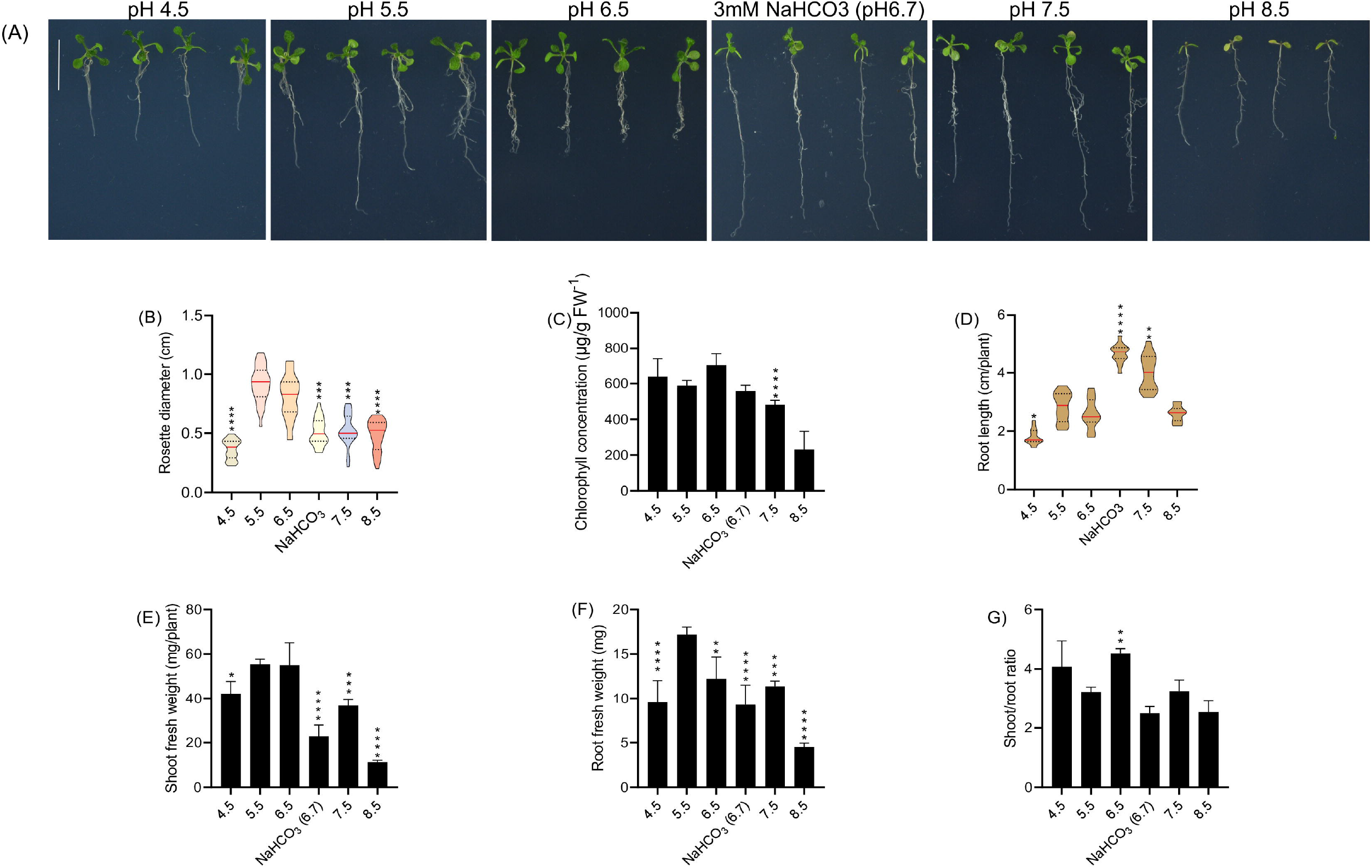
Phenotypic effects of varying media pH on plant growth. Plants were grown for 14 days under the various pH treatments. A, Phenotypes. B) Rosette diameter. C) Chlorophyll concentration. D), Root length. E, Shoot fresh weight. F), Root fresh weight. G) Shoot/root ratio. Values are mean ± SE of four replicates of four plants each. Statistical analysis was carried out using two-way Anova. No asterisk P > 0.05; *P ≤ 0.05; **P ≤ 0.01; ***P ≤ 0.001; **** P ≤ 0.0001.

### Exposure to high pH induces transport processes and tunes auxin homeostasis

To investigate the cause of growth cessation and leaf chlorosis of plants grown at alkaline pH, we transferred 10-day-old *Arabidopsis* Col-0 plants for six hours from slightly acidic (pH 5.5) to media adjusted to pH 7.5. Roots were subjected to gene expression analysis via RT-qPCR or transcriptomic analysis via RNA-seq. For the latter approach, a total of 42-62 million paired-end reads with a length of 150 base pairs were acquired for each library using the Illumina NOVAseq 6000 sequencing system and aligned to the Araport 11 annotation of the *Arabidopsis* genome. In total, 419 genes were differentially expressed between plants grown on pH 5.5 media and media adjusted to pH 7.5. Gene ontology categorization of the differentially expressed genes highlights redox reactions, phenylpropanoid/lignin metabolism, response to salicylic acid, and fatty acid metabolism as overrepresented categories in this data set (Supplementary Figure S1).

Unexpectedly, exposure to alkalinity strongly induced a suite of high-affinity nitrate transporters (i.e., *NRT2.6., NRT2.1, NRT2.4, NRT3.1* (*WR3*), and *NRT1.5*; Fig. 2; Supplementary Table S2). Since NRT transporters are NO_3_^−^/H^+^ cotransporters, the increased transcript levels are indicative of enhanced proton flux across the plasma membrane. Fitting this concept, also the expression of the rhizodermal urea/proton symporter *DUR3* (Kojima et al., 2007) was induced by elevated media pH (Fig. 2). *NRT2.1* (Δ RPKM ~800) and *WR3* (Δ RPKM ~400), suggested to function as a NRT2.1-WR3 hetero-oligomer (Yong et al., 2010), were the largest contributors to the increase in anion/H^+^-co-transport in terms of transcript levels (Fig. 2). Since plants were grown on media supplemented with adequate nitrate levels, induction of nitrate transporters does not seem to indicate a low N-status of the plants. Rather, this observation suggests a massive enrichment of protons in the cytosol, a supposition that is corroborated by the observation that the mitochondrial (proton-coupled) phosphate transporter *PHT3.2* was among the most strongly downregulated genes after transfer to alkaline media (Fig. 2). The proton/phosphate ratio for transporters of this type was estimated to be 2-4 (Młodzińska and Zboińska, 2016), suggesting rapid depletion of protons in the cytosol when this transporter is active.

**Figure 2.**
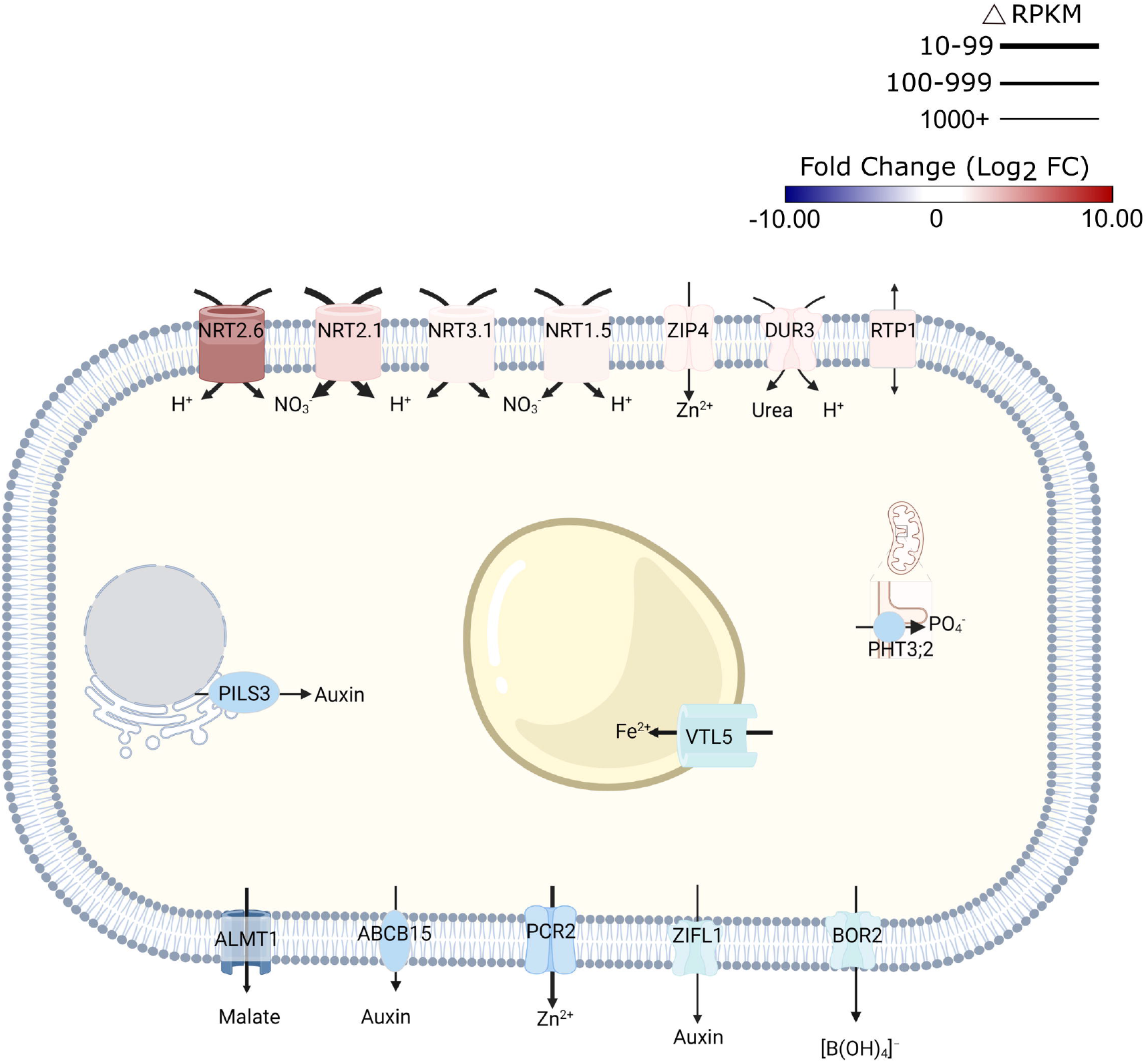
Schematic representation of transporters that are differentially expressed in response to short-term exposure to alkaline pH. Line thickness represents ΔRPKM values; colors represent log2FC. Arrows indicate the direction of transport. Figure was created with BioRender.com.

Expression of the malate transporter *ALMT1* was reduced in response to high pH (Fig. 2). Lower transcript abundance after transfer to pH 7.5 medium was also observed for the vacuolar iron transporter *VTL5*, presumably to avoid depletion of cytoplasmic iron. In addition to intracellular iron trafficking, uptake of Zn^2+^ from the soil solution via *ZIP4* was promoted at high pH; expression of the Zn^2+^ efflux protein *PCR2* was downregulated (Fig. 2).

Consistent with the assumption that elevated pH is compromising growth, several genes involved in auxin signaling and transport were found to be affected by transfer to high pH media. This subset includes several – mostly upregulated - small auxin up-regulated RNA (SAUR) genes (i.e., *SAUR38, SAUR40*, *SAUR41*, *SAUR44*, *SAUR55*, and *SAUR64*), likely to facilitate cell expansion. A function specifically in this process has been described for SAUR40 and SAUR41 (Qui et al., 2020). In addition, intracellular auxin homeostasis is affected through pH-dependent regulation of the auxin efflux genes *PILS3* and *ZIFL1* (Fig. 2; Supplementary Table S2). Together, these observations suggest that transmembrane proton transport, metal transport, and auxin homeostasis are the processes that are most prominently reflected in transcriptomic profile changes upon exposure to high pH.

### The expression of genes responsive to alkalinity is altered over a wide range of pH values

To obtain a more detailed picture on the pH-dependence of transcription, mRNA levels of selected genes from plants subjected to either short-term (6 hours) or long-term (14 days) growth on media with varied pH were determined by RT-qPCR (Fig. 3). Gene expression appears to be highly responsive to short-term changes in external pH over a range of pH values to which plants are realistically exposed in their natural habitats. The nitrate transporters *NRT2.4*, *NRT2.6* and *NRT3.1*, the plant intracellular Ras-group-related LRR *PIRL8*, the oxygenase *S8H*, and the DUF642 protein At2g41810 responded to the treatment with a continuous increase in transcript levels from low to high pH. A similar pattern was also observed for *EXPA17*, encoding an expansin that promotes lateral root formation (Lee and Kim, 2013), albeit expression of the gene was dramatically reduced at the highest pH. For another suite of genes comprising the laccase *LAC7*, the cytochrome P450 protein *CYP82C4*, the pathogen defense gene *MLO6*, the malate efflux transporter *ALMT1*, and the kinase *STRESS INDUCED FACTOR 1* (*SIF1*) an opposite pattern was recorded. *NRT1.1*, the main root nitrate transporter, showed a pronounced upregulation at low pH and a steep decrease in transcript levels at higher pH that plateaued at pH 5.5. Two of the tested genes, *SWEET12* and *SAUR44*, showed a more complex pattern (Fig. 3). Gene expression patterns observed after 6 hours remained largely unchanged when plants were exposed to the various media for an extended experimental period of 14 days (Supplemental Figure S2). It should also be noted that the bicarbonate treatment resulted in gene expression changes that were in line with the pH of the media and not altered by the presence of the salt as such.

**Figure 3.**
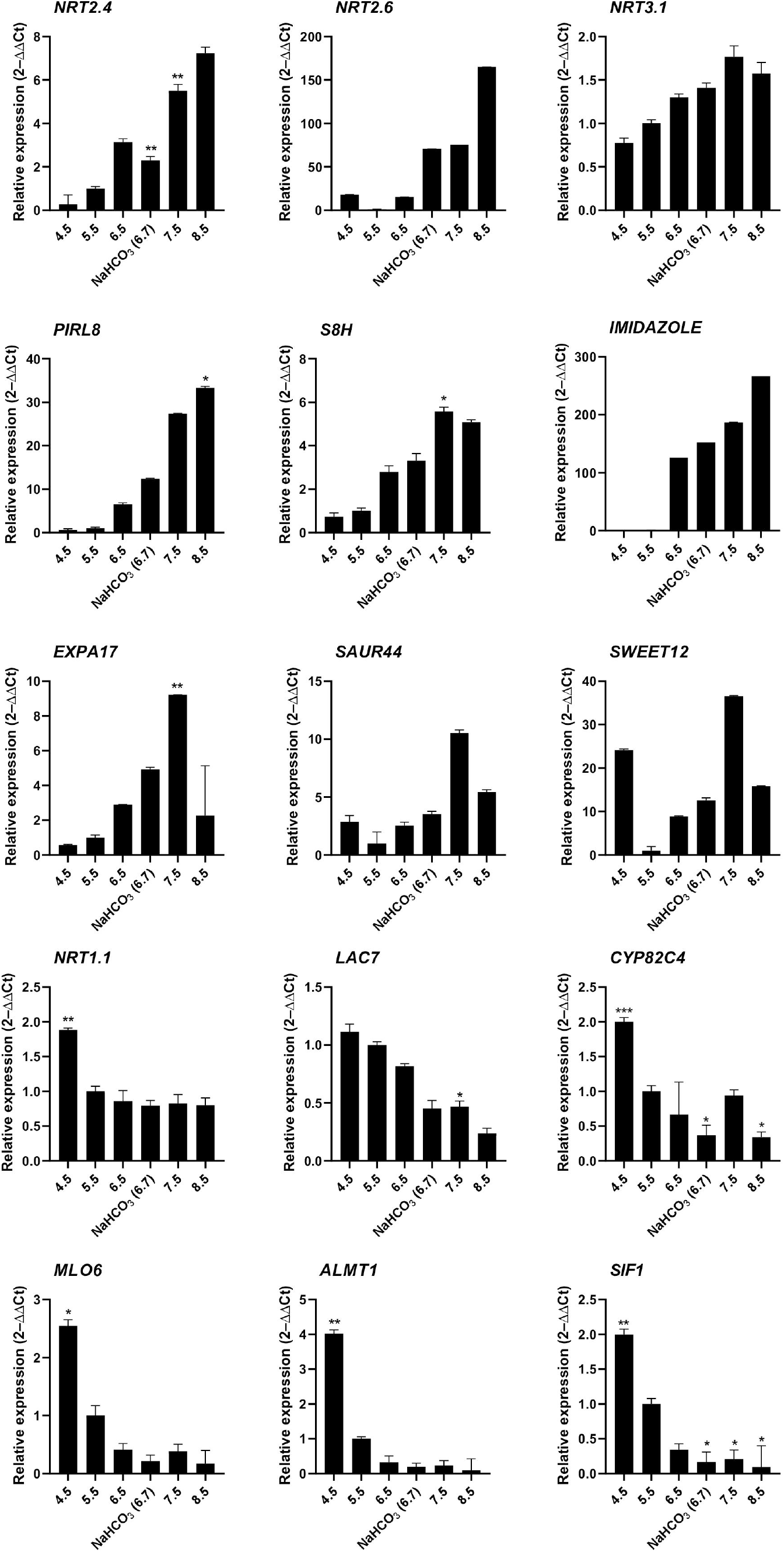
RT-qPCR analysis of the expression changes of selected genes in response to various pH treatments. Ten-day-old plants were transferred to solid media of varying pH for 6 hours. Gene expression values are normalized to EF1α and β-tubulin and calculated using the 2^−ΔΔCt^ method. Values are means □±□ standard deviation (SD) (n□=□3). No asterisk P = 0.05; * P ≤ 0.05; ** P ≤ 0.01.

### A comparison of the responses to alkalinity, iron deficiency, bicarbonate, and acidity identifies conserved gene clusters

In soils, elevated pH is generally associated with low availability of iron, excluding species with inefficient iron uptake systems from such habitats. To investigate whether exposure to elevated pH affects the iron status of the plants, we compared the transcriptome of pH 7.5 plants with the ‘ferrome’, a suite of genes that was found to be robustly modulated by iron supply across several studies (Hsieh et al., 2022). Whereas elevated pH strongly restricts the availability of iron *in situ*, in the current experimental set up iron mobility was sustained over a wide pH range by providing plants with the highly stable iron form FeEDDHA. A small subset of the ferrome genes was distincty affected in roots of plants exposed to pH 7.5, possibly in anticipation of the iron shortage generally associated with increased pH (Fig. 4). Pronounced upregulation was observed for the scopoletin hydroxylase *S8H*, a gene involved in the production of the iron-mobilizing coumarin fraxetin (Tsai et al., 2018; Gautam et al., 2021). Moreover, the vacuolar metal transporters *MTP8*, *IREG2* and *VTL5*, the clade Ib transcription factor *bHLH100*, *LAC7*, and the H_2_O_2_-response gene *HRG1* were affected by the pH 7.5 treatment (Gong et al. 2021). Interestingly, a suite of 17 (mostly downregulated) ferrome genes was also affected by low pH (Fig. 4), underlining the link between pH and iron availability. *HRG1* is the sole ferrome gene that is part of both the low Fe and high pH data sets.

**Figure 4.**
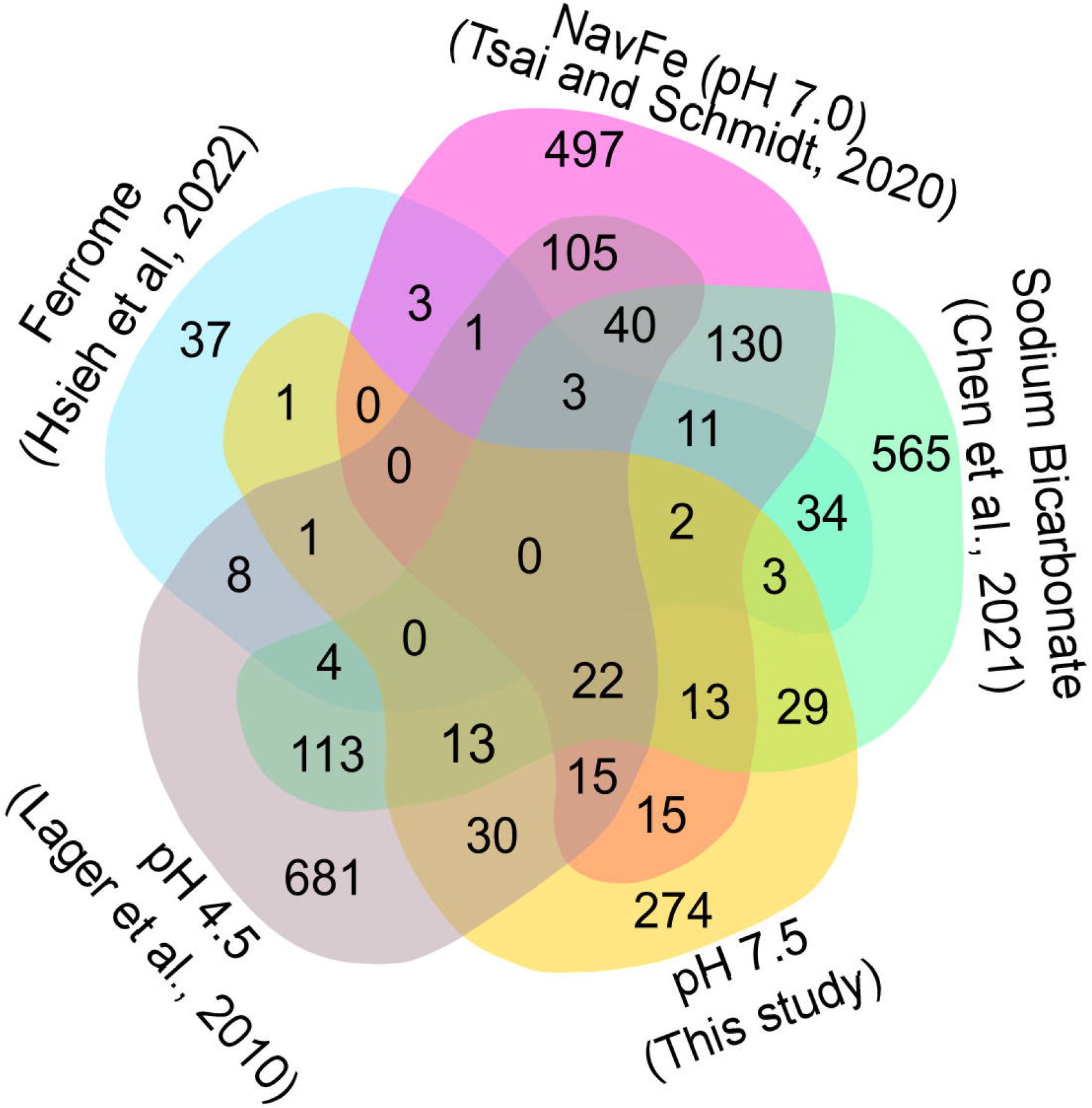
Venn diagram showing the commonalities of differentially expressed genes inferred from various RNA-seq derived data sets. Ferrome data are derived from Hsieh et al. (2022); NavFe data from Tsai and Schmidt (2020); data for bicarbonate-treated plants are taken from Chen et al. (2021) and normalized; pH 4.5 data are from Lager et al. (2010); data of pH 7.5 plants were generated in this study. The data were processed by trimming those that were statistically significant (*P* > 0.05) and differentially expressed with a log2 FC |1|.

The presence of high bicarbonate levels adversely affects the uptake of essential mineral nutrients, including iron (Chen et al., 2021). To investigate how the response to bicarbonate relates to that to iron deficiency and high pH as such, we compared the pH 7.5 data set with previously published transcriptomic changes in roots from plants grown on bicarbonate-supplemented media (Chen et al., 2021). Bicarbonate-treated plants mounted a pronounced iron deficiency response, affecting the expression of a subset of 47 ferrome genes (44%; Fig. 4). In addition, in this growth type a subset of 119 from genes responsive to pH 7.5 was differentially expressed. The relatively large overlap of genes affected by bicarbonate with the pH 7.5 and iron deficiency transcriptomes suggests that bicarbonate-treated plants strongly respond to both high pH and iron deficiency in addition to the responses triggered by the presence of bicarbonate as such.

We further compared the pH 7.5 data set with the transcriptome of plants grown on circumneutral media supplemented with an immobile iron source (non-available iron; navFe). Only a relatively small subset of the ferrome genes was among the genes that were differentially expressed between plants grown on navFe media (pH 7.0) and those grown on iron-deplete media at acidic pH (5.5), but a much larger set of genes (41) was comprised in the pH 7.5 transcriptome (Fig. 4). It thus appears that the response to high pH is substantially different from but partially overlapping with that observed in iron-deficient plants.

Since the expression of many pH 7.5 genes is also responsive to low pH, we included a data set derived from roots of plants exposed for 1 or 8 hours to pH 4.5 in the comparison (Lager et al., 2010). The transcriptomes of pH 7.5 and navFe plants showed a comparable percentage of genes responsive to low pH (19% and 21%, respectively; Fig. 4). This number was lower for bicarbonate-treated (12%) and for iron-deficient plants (14%), indicating that in these growth types the response to iron deficiency and bicarbonate is dominant and the overall transcriptional changes are less affected by changes in media pH.

### Genes involved in growth, pH homeostasis, and defense adapt plants to alkalinity

To identify key players conferring calcicole behavior, we compared differentially expressed genes within the five data sets with stringent thresholds (3-fold change on a log2 basis) and under the condition that a gene is differentially regulated in at least three out of the five conditions under study (Fig. 5). This conservative comparison highlighted three clusters of genes based on their susceptibility to a given suite of environmental conditions. The first group (cluster 1; ‘cell elongation’) comprises genes that were differentially expressed in all four pH-related treatments under investigation (i.e., pH 7.5, pH 4.5, navFe, and bicarbonate-treated plants). Three upregulated genes in this cluster encode proteins containing DUF642 domains. This domain is restricted to spermatophytes and putatively involved in homogalacturan esterification, a process associated with cell wall loosening. Two other upregulated genes in this group, the pectin-lyase At243890 and the expansin *EXPA17* also mediate cell wall-related processes. In addition, this cluster contains several genes involved in auxin homeostasis such as *SAUR41*, *SAUR44*, and the auxin transporter *PILS3*. Generally, genes affected by low pH were inversely regulated to those of the three high pH treatments. The cell elongation cluster also contains genes encoding putative signaling proteins or transcription factors that are largely undocumented such as *SIF1* and *PLANT INTRACELLULAR RAS GROUP-RELATED LRR 8* (*PIRL8*), representing the most dramatically regulated genes in this group. Notably, *PIRL8* was shown to be co-expressed with *EXPA17* and suggested to play a role in auxin-mediated lateral root growth (Hossain et al., 2022). *SENSITIVE TO PROTON RHIZOTOXICITY2* (*STOP2*), a homolog of *STOP1*, encodes the only transcription factor in this module.

**Figure 5.**
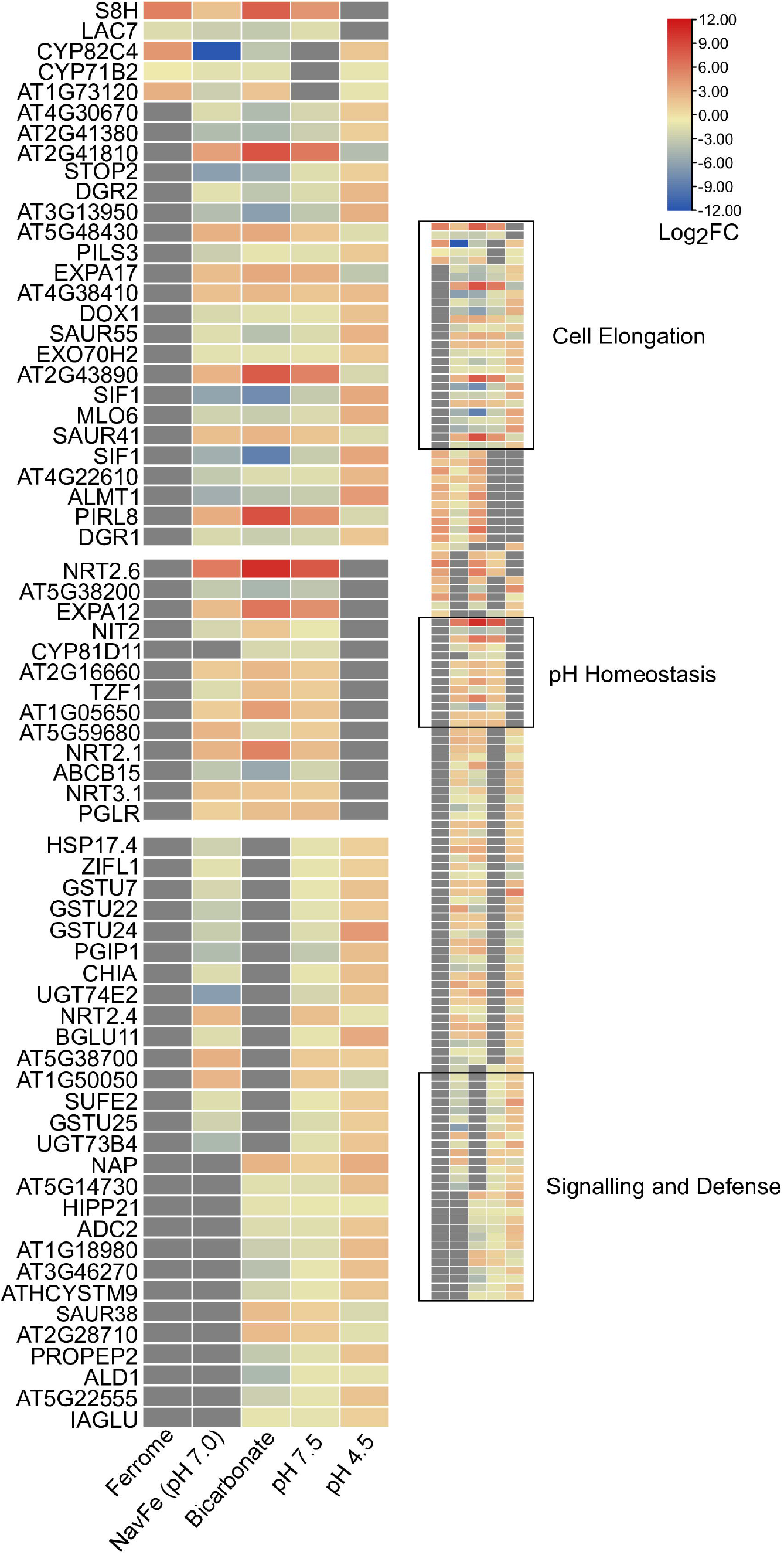
Heat map depicting genes identified in the co-expression networks. Colors representing log2FC range from red (upregulated) to blue (downregulated) as shown in the key; missing values shown in gray. Ferrome data are derived from Hsieh et al. (2022); NavFe data from Tsai and Schmidt (2020); data for bicarbonate-treated plants were taken from Chen et al. (2021) and normalized; pH 4.5 data are from Lager et al. (2010); data of pH 7.5 plants were generated in this study. The data were processed by trimming those that were statistically significant (*P*> 0.05) and differentially expressed with a log2FC |3|. Only genes that were differentially expressed in three or more treatments are included in the heatmap.

A second group (cluster 2; ‘pH homeostasis’) is comprised of genes that are solely regulated by treatments imposing elevated pH. Here, the two most strongly regulated genes within the pH 7.5 transcriptome are found, the nitrate transporter *NRT2.6* (up) and the nitrate-inducible class I glutamine amidotransferase-like superfamily protein At5g38200 (down). Furthermore, *NRT2.1*, *NRT3.1*, genes involved in cell wall modification, and an extra-cytoplasmatic LRR kinase (At5g59680) are part of this cluster.

The third module (cluster 3; ‘sensing and defense’) is comprised of genes responsive to both high and low pH (but not to iron) and contains two immune response-inducing secreted peptides, the CAP superfamily protein At1g50050, and the plant elicitor peptide *PEP2*. PEP1, a homolog of PEP2, was recently shown to sense extracellular alkalinity to induce immunity through binding to the receptor kinase PEPR (Liu et al., 2022a). The module also contains the so-far undocumented extra-cytoplasmatic receptor kinase At3g46270, which we named pHRK1 (pH-RECEPTOR KINASE 1). *pHRK1* and its homolog *pHRK2* (At3g46280) were both upregulated by elevated pH and previously found to be deregulated in *stop1* mutants (Sawaki et al., 2009).

### Co-expression analysis uncovers a putative pH homeostasis module

To further elucidate the processes that acclimate plants to external pH, we constructed networks based on pair-wise comparison of the co-expression relationships of differentially expressed genes using a data base of a total of 1,194 root-specific normalized RNA-seq data sets from the Sequence Read Archive hosted by the National Center for Biotechnology Information and the in-house developed MACCU toolbox (Lin et al., 2011). For pH 7.5 plants, the co-expression network features a subcluster comprising *pHRK1*, *STOP2*, *SIF1*, *LAC7*, *ALMT1*, *SAUR55*, and the alpha-dioxygenase *DOX1*(Fig. 6a). This sub-cluster appears to be conserved at low pH, but genes are generally regulated in the opposite direction (Fig. 6b; Fig. 7). Notably, this sub-cluster is also part of the co-expression network derived from bicarbonate-treated plants (Supplementary Fig. S3). Some of the genes observed in the network of pH 7.5 plants (i.e., *STOP2*, *EXPA17*, *LAC7*, *DOX1*, and *ALMT1*) are conserved in the network derived from navFe plants (Supplementary Fig. S4). The co-expression analysis suggests that a suite of co-expressed and putatively co-regulated set of genes is involved in the adaptation to external pH over a wide range of proton concentrations.

**Figure 6.**
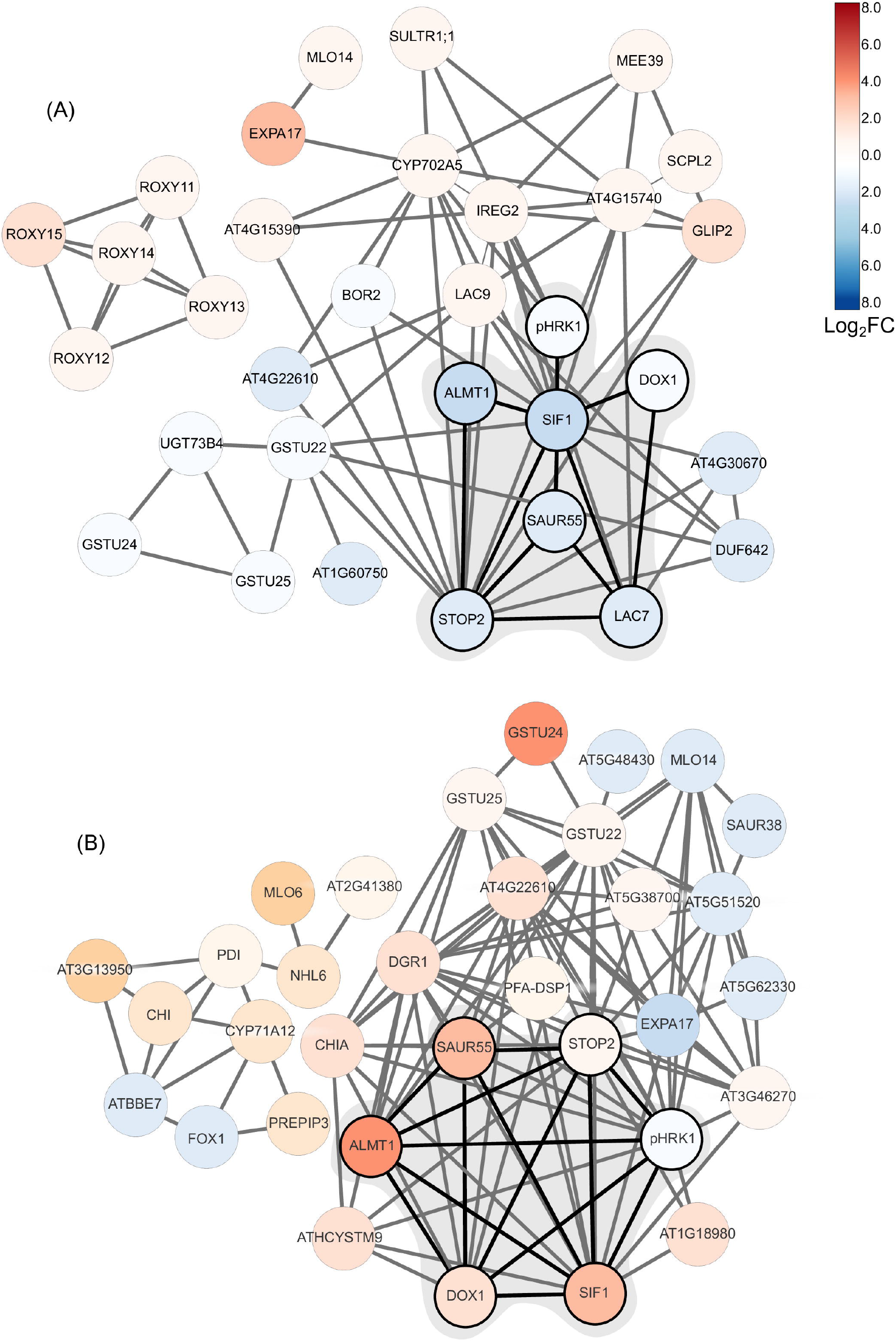
Co-expression network of DEGs from plants subjected to short-term changes in media pH. A) pH 7.5 plants (this study); B) plants subjected to pH 4.5 (Lager et al., 2010). Networks were generated using the in-house MACCU software package (Lin et al. 2011) against a data base comprising a suite of 1,194 normalized root-related RNA-seq data sets. Networks were constructed using a Pearson’s correlation coefficient of <0.8 for pairwise expression of genes that were differentially expressed upon exposure to pH 7.5 or pH 4.5. Networks were drawn using Cytoscape. Log2C values are color coded according to the key.

**Figure 7.**
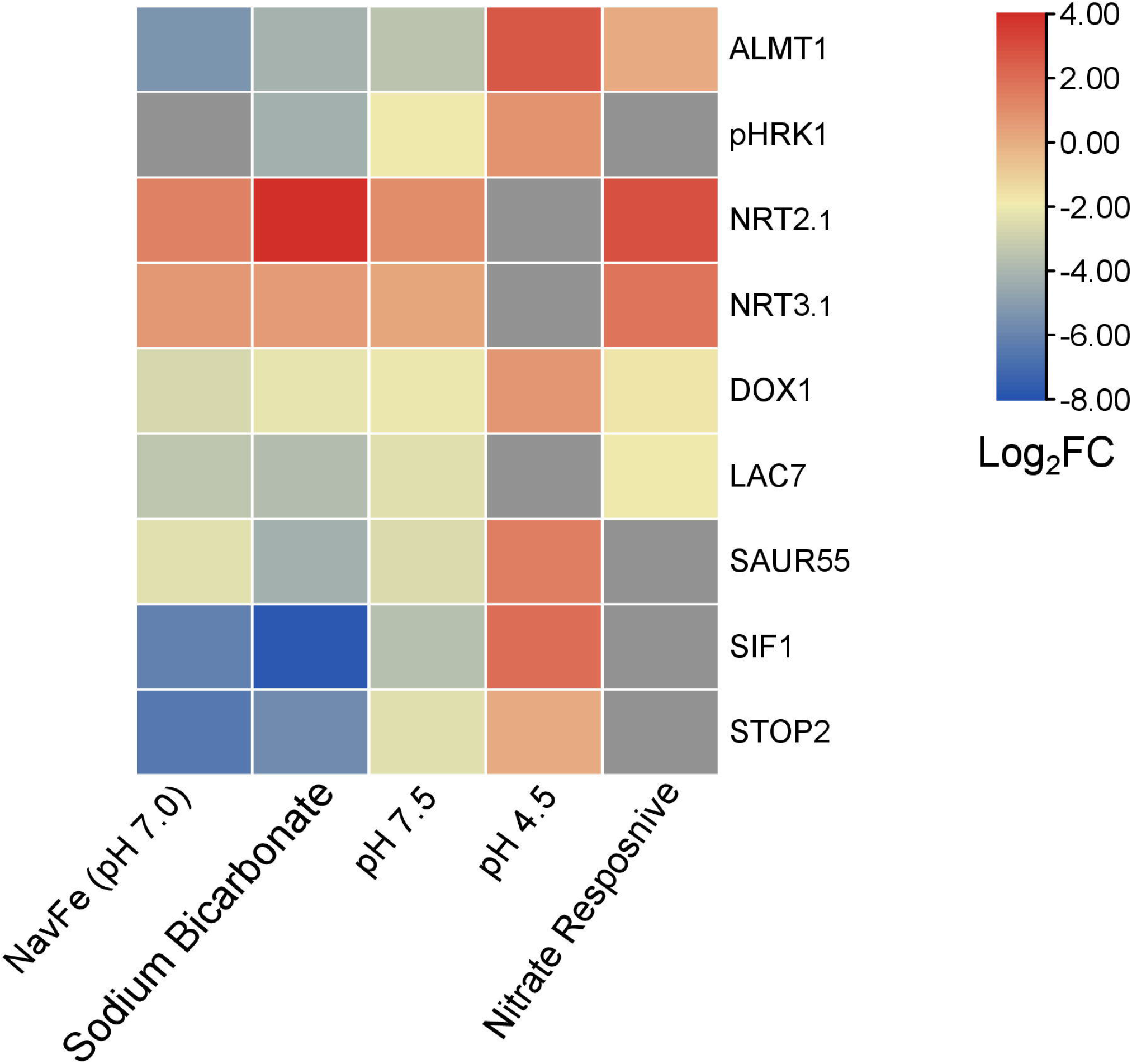
Heat map depicting genes that are differentially expressed in various treatments. NavFe data from Tsai and Schmidt (2020); data for bicarbonate-treated plants were taken from Chen et al. (2021) and normalized; pH 4.5 data are from Lager et al. (2010); nitrate-responsive data are from Vidal et al. (2013); data of pH 7.5 plants were generated in this study. The data was processed by trimming those that were statistically significant (P > 0.05) and differentially expressed with a log2FC |2|. Only genes that were differentially expressed in three or more treatments were included in the heatmap.

## Discussion

### Bicarbonate, but not alkalinity as such triggers an iron deficiency response

Similar to what has been observed for plants responding to low pH (Lager et al., 2010), we show here that short-term exposure to moderate alkalinity leads to pronounced changes in the transcriptomic profile of *Arabidopsis* root cells. The phyto-availability of iron is closely related to soil pH and decreases dramatically at elevated pH. It can thus be speculated that alkalinity induces a response to iron deficiency, either as a consequence of ‘true’ iron depletion within the plant, caused by impaired uptake of iron, or in anticipation of the lower iron supply generally associated with alkaline substrates. This does, however, not seem to be the case; only a minor fraction of the ferrome genes was affected by raising the media pH. The small overlap of the two data sets does not seem to be caused by the short experimental period; a six-hour-transfer to iron-deplete media was sufficient to induce a much more pronounced iron stress response (Hsieh et al., 2022). The supposition that high pH and iron deficiency trigger separate signaling cascades is in line with the observation that restricting iron availability at circumneutral pH (navFe plants) does not simply intensify the expression of ferrome genes (Tsai and Schmidt, 2021). Rather, such treatment induced a separate, largely pH-dependent set of genes.

Long-term bicarbonate-treated plants (Chen et al., 2021) showed a more pronounced response to both iron deficiency and alkalinity than pH 7.5 plants, albeit the pH change induced by bicarbonate was less severe. Notably, short-term exposure to bicarbonate exerted a response that was not different from the influence of pH as such (this study). However, when we exposed plants to bicarbonate for a 14-day-period, growth was significantly more restricted than expected from the effect on pH, suggesting that the presence of bicarbonate enforces other constraints to growth and the homeostasis of mineral nutrients such as iron. The effect of bicarbonate on pH- and iron-responsive genes might be aggravated by the strong buffering effect of the salt, compromising compensatory measures to recalibrate both apoplastic and cytoplasmic pH, as well as iron distribution and homeostasis.

### External pH alters proton fluxes through modulation of anion/H^+^ co-transporters

The free cytosolic H^+^ concentration is in the sub-micromolar range (Wegner et al., 2021), and must be carefully balanced to avoid fluctuations in cytosolic pH and subsequent perturbations of cellular functions. Adjustment of cellular pH is thought to be mediated by a combination of H^+^-producing and H^+^-consuming reactions in concert with H^+^ movements across the plasma membrane (Smith and Raven, 1979). Changes in the activity of NO_3_^−^/H^+^ co-transporters significantly affect the pH of both the apoplast and the cytosol (Fan et a., 2016; Zhou et al., 2021). The current analysis suggests that dramatic changes in anion-coupled transmembrane H^+^-fluxes are a general feature of plants exposed to alkaline pH; increased expression of nitrate transporters was observed in pH 7.5, navFe, and bicarbonate-treated plants. Of note, induction of NRT homologs in response to alkaline pH was observed in a proteomic survey of sugar beet plants (Geng et al., 2021), suggesting that this response could be conserved across species. Upregulation of *NRT2.1* and *NRT3.1* was also reported for nitrate-treated plants (Vidal et al. 2013), a circumstance that was associated with the induction of enzymes related to the assimilation of nitrate such as the nitrate reductases NR1 and NR2. Conspicuously, in all three high pH treatments, the increase in transporter abundance does not seem to be correlated with nitrate assimilation. *NR* genes were either not regulated (in pH 7.5 and navFe plants) or dramatically repressed (in bicarbonate-treated plants) upon exposure to elevated pH (Supplementary Table S3). For example, *NR1* was 3-fold upregulated in nitrate-starved plants, but 5-fold downregulated upon exposure to bicarbonate. This observation resembles the regulation of other genes encoding enzymes involved in nitrate assimilation such as the glutamine synthetases *GLN2* and *GSR2* as well as the glutamate synthase *GLT1*, which are inversely regulated in nitrate-starved and bicarbonate-treated plants (Supplemental Table S3). Similar to pH 7.5 plants, the glutamine amidotransferase At5g38200 was strongly downregulated in bicarbonate plants but upregulated in nitrate-treated plants. It thus appears that the uptake, but not the assimilation of nitrate uptake is upregulated by high pH.

The supposition that proton homeostasis is the driving force behind the induction of nitrate transporters is supported by the pronounced upregulation of other anion/H^+^ co-transporters such as *DUR2* and *NRT2.6*, a common observation across all high pH treatments. Further support for this assumption comes from the downregulation of intracellular anion/H^+^ co-transporters such as *PHT3;2* (in pH 7.5 plants) and *SULTR3.5* (in bicarbonate-treated plants), which mediate transport processes that would deplete protons in the cytoplasm. *PHT3;2* and *SULTR3.5* are massively up-regulated in nitrate-starved plants, further supporting the idea that nitrate is triggering a response distinct from that to high pH. Also of note, expression of the nitrate exporter *NAXT1* was repressed in both nitrate-starved and bicarbonate-treated plants. NAXT1 protein abundance is induced by strong acidosis (Segonzac et al., 2007), further indicating that nitrate homeostasis is differently affected by external pH.

### Acidic and alkaline pH induce distinct nitrate transporters

The nitrate transporter NRT1.1 is recruited to recalibrate apoplastic pH when plants are exposed to acidity (Ye et al., 2021). In line with this concept, our data confirm that acidic pH supports *NRT1.1* expression, an observation that has been reported earlier (Tsay et al., 1993). Notably, NRT1.1 activity was shown to impair the utilization of iron in leaves by alkalization of the apoplast (Ye et al., 2022). Since iron availability is severely restricted in an alkaline environment, a decrease rather than an increase in proton-coupled co-transport processes should be expected when plants are subjected to elevated pH. The rationale behind the – in terms of iron acquisition counterintuitive - induction of nitrate transporters upon exposure to both low and high pH may be simply a matter of priorities. Cellular pH homeostasis might be difficult to achieve at an external pH of 7.5, which may increase cytosolic pH and massively disturb cellular functions without intervention. At acidic pH, an increase in apoplastic pH is essential to avoid cell damage due to Al^3+^ toxicity, but the protons taken up alongside nitrate on cytosolic pH are not severely affecting the pH gradient across the plasma membrane since the external pH differs only by one pH unit from the optimal growth conditions for the calcifuge species *Arabidopsis thaliana*.

The answer to the question as to why different NO_3_^−^ transporters are recruited at low and high pH (i.e., NRT1 vs NRT2) may lie in the fact that this differential recruitment upon high pH have distinct effects on root architecture. NRT2.1 was shown to promote lateral root initiation (van Gelderen et al., 2021; Remans et al., 2006), supporting topsoil foraging of phosphate, a scarce resource in a high pH environment. The effect of NRT2.1 on root development is, however, highly dependent on the N-status of the plant (Little et al., 2005), and may produce a distinct readout in calcareous soils. In rice, overexpression of *OsNRT2.1* enhanced elongation of primary roots (Naz et al., 2019), an effect that could counteract root growth inhibition at alkaline pH. It is thus tempting to assume that the recruitment of different NRT transporters in response to the external pH can adapt root growth and architecture to the prevailing edaphic conditions. Crucially, in both pH 7.5 and bicarbonate-treated plants, a group of five nitrate-inducible thioredoxins (GRXS) were upregulated. Overexpression of *GRXS* genes was shown to suppress the expression of the nitrate transporters *NRT2.6* and *NRT3.1* (Ehrary et al., 2020). It can be assumed that *GRXS* genes are induced at high pH to balance the pronounced induction of genes involved in nitrate uptake in response to high pH.

### A conserved gene module orchestrates root growth in response to external pH

The acid growth theory postulates that high apoplastic pH reduces growth by stiffening the cell wall and compromising cell elongation. How root growth is tuned in response to soil pH has yet to be elucidated. The current survey highlights a suite of robustly pH-regulated genes comprising *pHRK1*, *SIF1*, *LAC7*, *DOX1, SAUR55, STOP2*, and *ALMT1* that may play a role in pH-dependent growth regulation. Of particular interest is the extracellular receptor kinase *pHRK1*, which is co-expressed with genes involved in iron uptake and homeostasis such as the putative iron sensors *BTSL1* and *BTSL2* (Supplemental Fig. S5). The exact link between high pH signaling and iron metabolism remains to be established. pHRK1 has predicted interactions with the GRAS family transcription factor protein At5g19970, a member of the ferrome (Supplemental Fig. S6). In line with such an interconnecting role, At5g19970 is responsive to iron deficiency, bicarbonate, nitrate, and navFe (although slightly exceeding the set P-value of 0.05 in the latter case). It thus appears that the responses to high pH, iron deficiency, and – to a certain degree – nitrate are interconnected by some promiscuous nodes that are orchestrating the responses to the plant to a given set of edaphic conditions (Fig. 7).

The pronounced downregulation of *ALMT1* in all high pH treatments was an unexpected observation. Since the function of ALMT1 is Al^3+^ detoxification via malate secretion (Hoekenga et al., 2006; Kobayashi et al., 2007), one would not expect much difference in transcript abundance of this transporter between plants grown at optimal (pH 5.5) and alkaline pH. ALTM1 is, however, also critical in regulating root meristem activity and, thus, root growth. In response to phosphate deficiency, aberrant iron precipitation in the apoplast mediated by the multi-copper oxidase LPR2 and peroxidase-mediated redox cycling of iron lead to meristem exhaustion, cell wall stiffening, and subsequent growth arrest (Müller et al., 2015; Balzergue et al., 2017). In this setting, phosphate deficiency promotes the expression of *ALMT1* and represses primary root growth. Similar to phosphate deficiency, NH_4_^+^ toxicity leads to inhibition of primary root growth (Rogato et al., 2010). A forward genetic screen for mutants that are insensitive to NH_4_^+^-induced root growth cessation identified LPR2 as being crucial for this process (Liu et al., 2022b). Similar to what has been observed under phosphate starvation, NH_4_^+^-induced root growth inhibition is associated with massive iron precipitation in the apoplast (here exclusively mediated by LPR2), which is upregulated by NH_4_^+^. NH_4_^+^-induced redox cycling of iron is associated with ROS formation, which in turn compromises sucrose transport in the phloem. Increasing NO_3_^−^ uptake prevents aberrant iron deposition, thereby sustaining phloem function and sucrose supply to root cells. Phloem integrity is crucial for sustained root growth to counteract the inhibitory effect of high external pH. In fact, in the current survey, the sucrose transporters *SWEET11* and *SWEET12* were induced at high pH. Expression of the multi-copper oxidases *LAC7* and *LPR1/LPR2* is highest in the stele (http://bar.utoronto.ca/eplant), reside in the extracellular matrix and have similarities regarding their physiochemical properties (Supplementary Figure S7). It is tempting to assume that *ALMT1*, *LAC7*, and *DOX1* tune growth in response to and in concert with external pH. In this scenario, LAC7 would modulate root meristem activity under normal and acidic growth conditions, while the gene is strongly repressed at high pH, where root growth is hampered by reduced cell wall extensibility.

Support for this concept comes from another observation. Two nitrate-inducible GARP-type (NIGT) transcriptional repressors involved the integration of N- and P-signaling, *HHO1* and *HRS1*, were found to be strongly downregulated in bicarbonate-treated plants. Phosphate deficiency induces the expression of *NIGT* genes, thereby inhibiting NRT2.1-mediated nitrate uptake and primary root growth (Medici et al., 2015). Notably, *HHO1* and *HRS1* are induced in nitrate-starved plants, possibly to regulate *NRT2.1* expression when N supply is restored. Together these data suggest that downregulation of the *LAC7*/*DOX1*/*ALMT1* module at high pH adapt root meristem activity to the prevailing external pH.

## Conclusions

The present survey allows us to infer a suite of adaptations that aid plants to thrive in alkaline substrates, comprising genes putatively involved in signaling extracellular pH and adjustment of intracellular pH. We further found that exposure to high pH modulates the transcript levels (as a proxy for altered activity) of anion/H^+^ coupled transport processes in a way that secures the enrichment of protons in the cell interior, possibly to counteract alkalization of the cytoplasm. The regulation of root architecture by nitrate and nitrate transporters is complex and interlinked with auxin transport (Jia and von Wirén, 2020), making predictions regarding such changes in response to high pH vague with the information at hand. However, the consistency and robustness of the changes in the expression of genes involved in proton-coupled transport processes strongly imply that these changes adapt plants to alkalinity. Lastly, we show that a robustly co-expressed module comprising *ALMT1*, *LAC7*, and *DOX1* suppositionally regulates primary root growth in a pH-dependent manner. While the physiological role and ecological relevance of these adaptive processes awaits experimental validation, we believe that our study provides guidance for follow up research that will help to elucidate the molecular basis of calcicole behavior.

## Author Contributions

MB, EJH, HHT, and AR conducted experiments and analyzed the data. WS conceived the project, analyzed the data and drafted the manuscript. All authors edited the manuscript and approved the final version of the draft.

## Acknowledgments

We thank Shou-Jen Chou from the Genomic Technology Core Facility at IPMB for preparing libraries for RNA-seq analysis. We also thank Mei-Jane Fang from the Genomic Technology Core Facility at the Institute of Plant and Microbial Biology for using the QuantStudio 12K Flex Real-Time PCR System. The authors are most grateful for the support provided by Dr Wendar Lin from the Bioinformatics Core Facility at IPMB, Academia Sinica. We acknowledge the use of the BioRender software (https://biorender.com) for generating the figures of this article.

## Supplementary Material

**Supplementary Table S1.** Primers for RTqPCR

**Supplementary Table S2.** Differentially expressed genes upon exposure to pH 7.5 media.

**Supplementary Table S3.** Regulation of genes encoding proteins involved in nitrate assimilation upon various treatments.

**Supplementary Figure S1.** Overrepresented GO categories in the pH 7.5 data set.

**Supplementary Figure S2.** RT-qPCR analysis of selected pH-dependent genes determined 14 days after exposure to the various treatments.

**Supplementary Figure S3.** Co-expression network of bicarbonate-treated plants.

**Supplementary Figure S4.** Co-expression network of navFe plants

**Supplementary Figure S5.** Co-expression network around pHRK1.

**Supplementary Figure S6.** PPI network of pHRK1.

**Supplementary Figure 9:** Sequence alignment of the multi-copper oxidases LPR1, LPR2, and LAC7.

## Data Availability Statement

The data sets for this study can be found at the NCBI depository. BioProject: PRJNA893419. SRA accessions: SRR22018204~9.

