## Supplementary material for "Alkalinity modulates a unique suite of genes to recalibrate growth and pH homeostasis": Table S1

**Supplemental Table 1. Table of qPCR primers for real time measurement assays.**

| Name | Gene ID | Forward (5' to 3') | Reverse (5' to 3') |
| --- | --- | --- | --- |
| NRT2.4 | AT5G60770 | TGGTCTTACTGGAGCTGGTG | GTTCCGTGGAGAACGTTGAG |
| NRT2.6 | AT3G45060 | AACCGACATCGGAAATGCTG | AAGGTCACATACAGCACCCA |
| NRT1.1 | AT1G12110 | GACTTCGACCACCAAGATGC | GAGCGGTGAGGTATAAGGCT |
| NRT3.1 | AT5G50200 | CGGCAAGGATACGTTGAACA | ATGGTCGGTCAACTTGGCTA |
| LAC7 | AT3G09220 | CTAACGGTTTGGGCAGATGG | TTGGCCGGTGATGTTGAATC |
| PIRL8 | AT4G26050 | CGCCAACTTCAACGAGCTTA | ACGGAGAGCTTCGTCAGATT |
| CYP82C4 | AT4G31940 | CTTACATGGGCCATTTCTC | TCCTCGACGTTCCTGTCTCT |
| S8H | AT3G12900 | GGTCATCAACATCGGCGACA | ACAACTTCCGGCAAGGGACC |
| MLO6 | AT1G61560 | ATGCTCCTCACAAACGAAGC | GAGACTCAGGATCCCACGAA |
| EXPA17 | AT4G01630 | ACACAGACGGCTACAAGACA | GGTAACATCCACCGCATGAC |
| SAUR44 | AT5G03310 | CACGATCCCATCTTCAGAGC | TCGTCACAAGCAATACAAAGC |
| Imidazole | AT2G41810 | CTCTCGTTCTCTCTCTTTGCTTGG | TCCCATTTGGGAGAAGCCCATC |
| SWEET12 | AT5G23660 | TCGGATTCTCTGTCTGCGTT | ACGGCATGTACTCCACACTT |
| ALMT1 | AT1G08430 | GCGATGAAGAAGCTTTGGCT | AATGCACCAACGGCAACATA |
| SIF1 | AT1G51840 | CGCCAAGGATTTCGAACCAT | TCGCGGGTGACATTTAGATTG |
| Tubulin | AT1G20010 | CGACAATGAAGCTCTCTACGA | AAGTCACACCGCTCATTGTT |
| Eflα | AT5G60390 | GAGCCCAAGTTTTTGAAGA | CTAACAGCGAAACGTCCCA |
