## Supplementary material for "Alkalinity modulates a unique suite of genes to recalibrate growth and pH homeostasis": Table S3

**Supplementary Table S3.** Regulation (log2FC) of genes encoding proteins involved in nitrate assimilation upon various treatments.

| Gene Name | AGI | Lager et al., 2010 | Chen et al., 2021 | Schmidt, This study | Tsai and Schmidt, 2020 | Vidal et al., 2013 |
| --- | --- | --- | --- | --- | --- | --- |
| NR2 | AT1G37130 | 1.12 | -2.53 |  | -1.47 | 1.34 |
| NR1 | AT1G77760 |  | -5.03 |  |  | 3.04 |
| NiR1 | AT2G15620 |  | -2.64 |  |  | 4.28 |
| Gln 1.5 | AT1G48470 |  |  |  |  |  |
| Gln 1.2 | AT1G66200 |  | -1.38 |  |  | 1.25 |
| Gln 1.3 | AT3G17820 |  | 1.24 |  |  |  |
| Gln 1.1 | AT5G37600 |  |  |  | 1.141 | -0.96 |
| GLN2 | AT5G35630 |  | -0.64 |  |  | 1.67 |
| SYNTHEA | AT5G38200 |  | -4.51 | -3.44 | -3.20 |  |
| GSR2 | AT1G66200 |  | -1.38 |  |  | 1.25 |
| Glu1 | AT5G04140 |  | -2.72 |  |  |  |
| Glu2 | AT2G41220 |  | -0.71 |  |  |  |
| GLT1 | AT5G53460 |  | -1.40 |  |  | 2.26 |
| GDH 1 | AT5G18170 | 1.78 | -1.06 |  |  |  |
| GDH 2 | AT5G07440 | 1.83 |  | -1.21 |  |  |
| GDH 3 | AT3G03910 |  |  |  |  |  |
| BT1 | AT5G63160 |  |  |  | -1.36 |  |
| BT2 | AT3G48360 |  |  |  | -1.46 | 3.46 |
| NAXT1 | AT3G45650 |  | -1.43 |  |  | -1.26 |
| NAC056 | AT3G15510 |  | 1.92 |  | -1.72 |  |
| ROXY2 | AT5G14070 |  |  |  | -3.96 |  |
| ROXY4 | AT3G62950 |  | 2.47 |  |  |  |
| ROXY5 | AT2G47870 |  | 1.81 |  |  |  |
| ROXY7 | AT2G30540 |  | 3.97 |  |  |  |
| ROXY11 | AT4G15700 |  | 3.14 | 1.27 |  |  |
| ROXY12 | AT4G15690 |  | 2.66 | 1.43 |  |  |
| ROXY13 | AT4G15680 |  | 2.88 | 1.25 |  |  |
| ROXY14 | AT4G15670 |  | 3.80 | 1.02 |  |  |
| ROXY15 | AT4G15660 |  | 2.71 | 1.85 |  |  |
| ROXY17 | AT3G62930 |  | 0.87 |  |  |  |
| ROXY18 | AT1G03850 |  | -0.80 |  | -1.97 |  |
| ROXY19 | AT1G28480 | 1.86 | -1.97 |  |  |  |
| ROXY21 | AT4G33040 |  | -1.12 |  |  | -2.33 |
