## Supplementary figures and images for "Alkalinity modulates a unique suite of genes to recalibrate growth and pH homeostasis"

### Figure S1

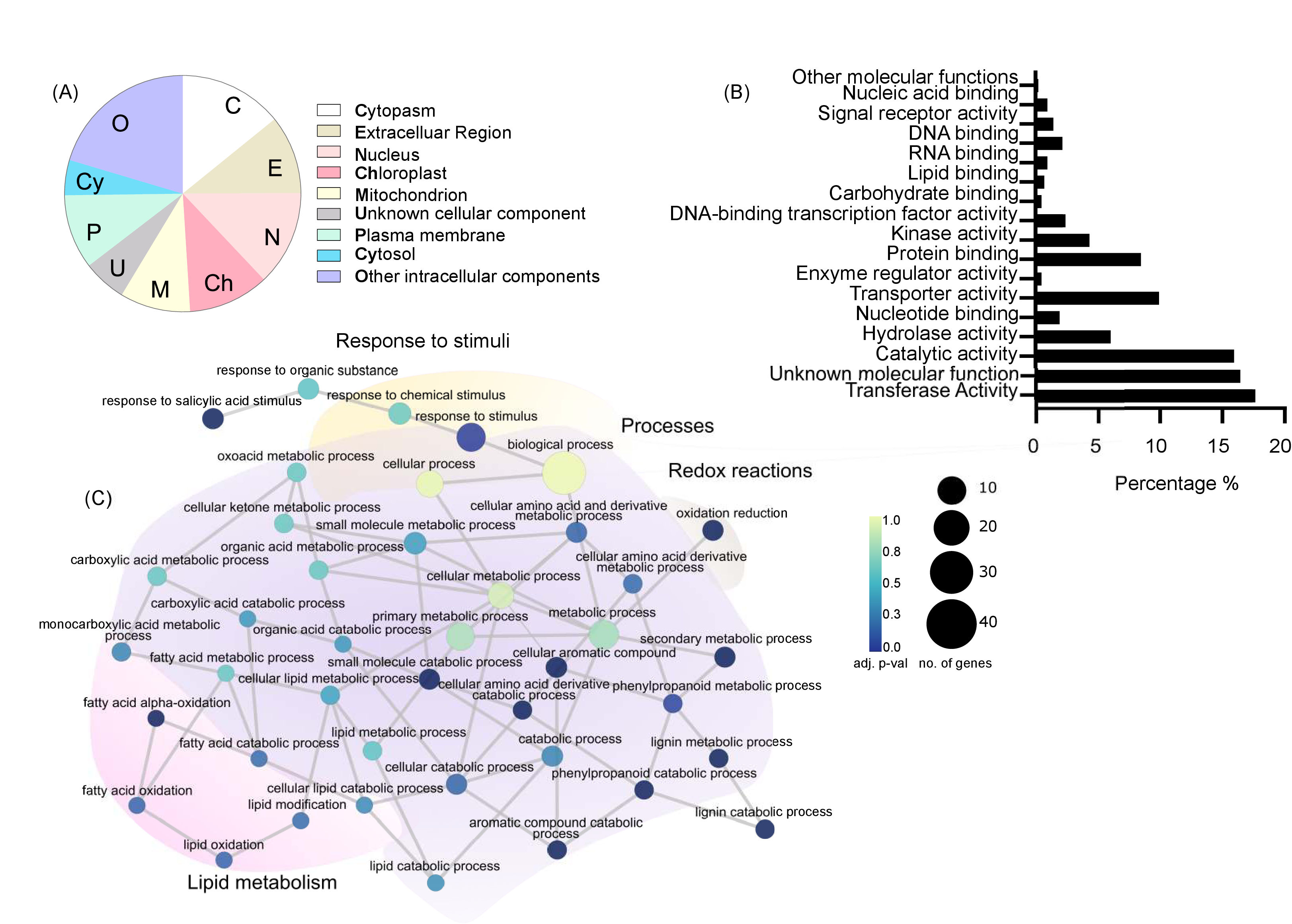

### Figure S2

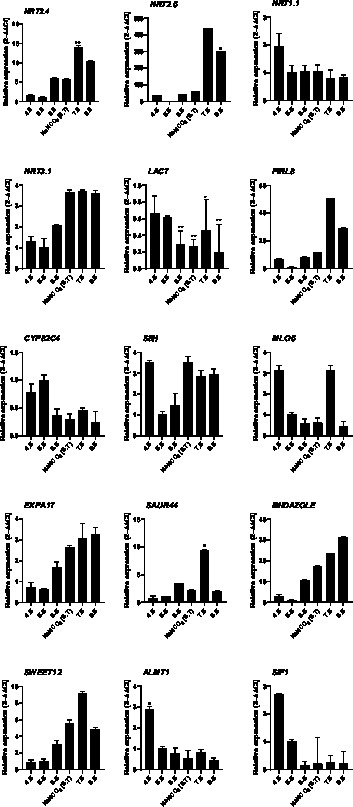

### Figure S3

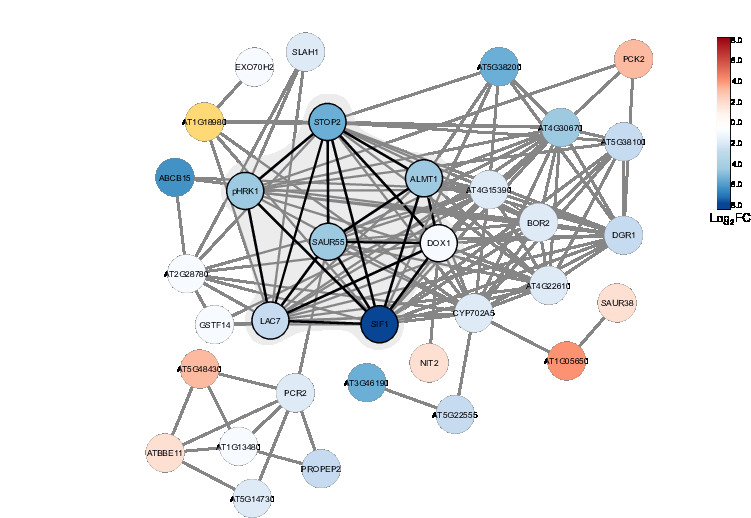

### Figure S4

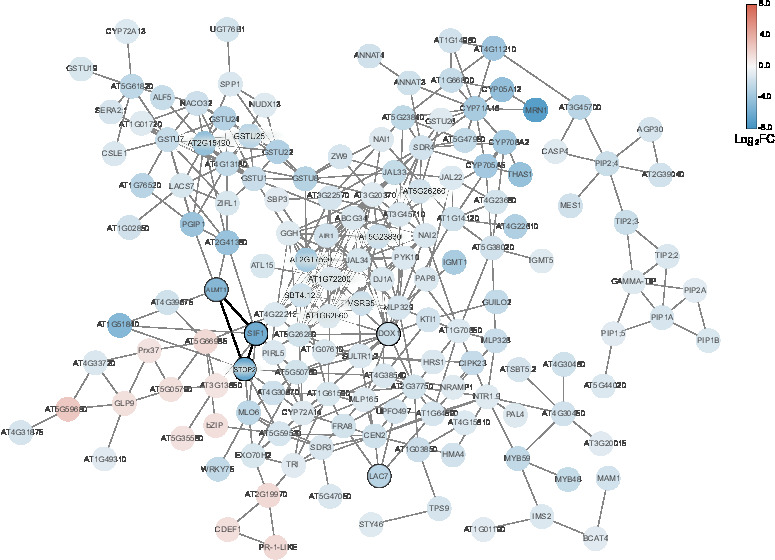

### Figure S5

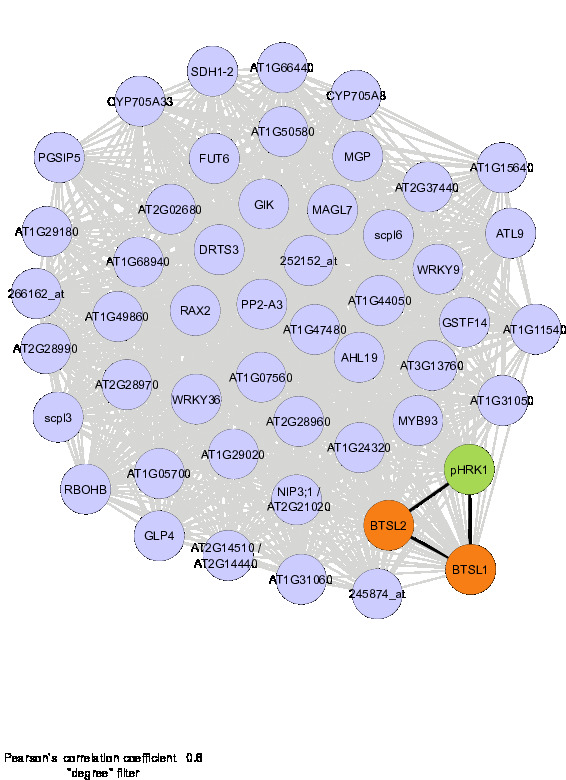

### Figure S6

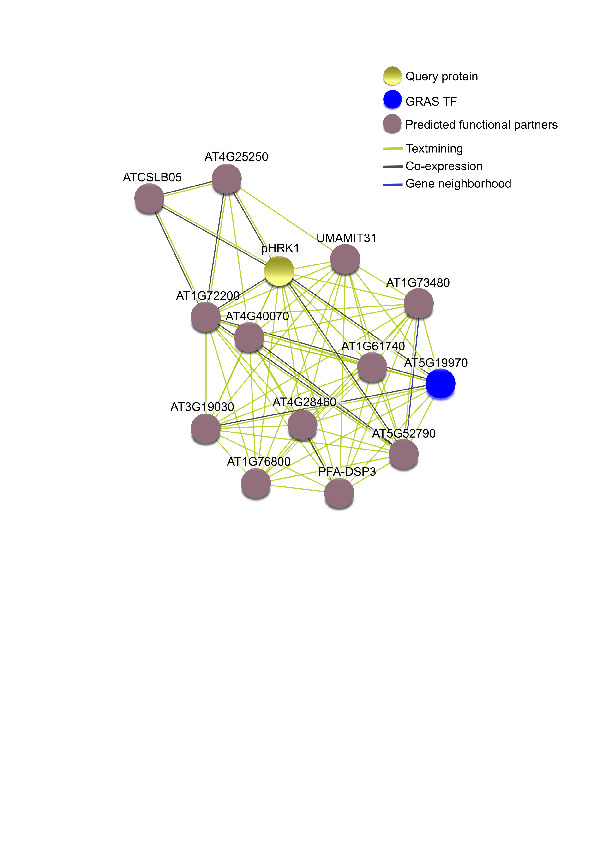
